# Radiosensitizers in Cancer Therapy: A Global Bibliometric Analysis

**DOI:** 10.1101/2025.06.26.661878

**Authors:** Shreyas Annagiri, Paul M. Harary, Yusuke S. Hori, Fred C. Lam, Deyaldeen AbuReesh, Armine Tayag, Lousia Ustrzynski, Sara C. Emrich, Steven D. Chang, David J. Park

## Abstract

**Background/Objectives:** Radiosensitizers are compounds given concurrently with radiotherapy to enhance the killing of cancer cells while sparing healthy tissue. Radiosensitization research has involved a diverse range of therapeutic agents. In the present study, we aimed to investigate international trends in the development of radiosensitizers across the most common cancer types.

**Methods:** A bibliometric analysis was performed of the field of radiosensitizer research from 1956 to 2024. Individual author impacts and trends, collaborations between and productivity of countries, and themes/keywords were analyzed.

**Results:** Our search yielded 12,690 results. The most highly represented countries were the United States of America, China, and Germany. Radiosensitizer studies for breast cancers demonstrated the highest rate of annual growth in record count by comparison with other cancer types, while publications for gynecological cancers showed the slowest growth. The most common radiosensitizers investigated included ATM kinase inhibitors, chemotherapies, gold nanoparticles, mTOR inhibitors, natural compounds such as caffeine or curcumin, and poly (ADP-ribose) polymerase inhibitors.

**Conclusions:** The United States, Germany, and China were the most productive countries during the study period, with China demonstrating the greatest increase in annual publication rate. Additionally, pre-clinical studies primarily investigated gold particles and targeted therapies. By comparison, clinical studies focused on radiosensitizing chemotherapies.

## Introduction

Radiation therapy represents a key treatment modality for many cancers, both in the standalone setting and in combination with chemotherapy or surgery. A large range of techniques exist for radiation delivery, with a growing role for conformal and image-guided approaches (1). Radiation causes cell death and DNA damage through both the direct destruction of genetic material from ionizing radiation and the generation of reactive oxygen species (ROS) which induce single and double-stranded DNA breaks (2,3). However, cancers may develop resistance to radiotherapy due to factors such as a hypoxic tumor microenvironment, angiogenesis, alterations in metabolism, proliferation of cancer stem cells, and intratumoral heterogeneity. To overcome this radioresistance and enhance the therapeutic index of treatment, radiosensitizers may be given concurrently with radiotherapy. These are therapeutic agents which may potentiate the killing of cancer cells while sparing healthy tissue (4). Mechanisms of radiosensitization are diverse and include the creation of cytotoxic substances (5), inhibition of repair mechanisms (6), targeting of hypoxic cells (7), or a combination of approaches (8). For instance, gold nanoparticles act as radiosensitizing agents by facilitating the absorption of greater amounts of radiation and enhancing the production of secondary electrons and ROS to damage cancer cell DNA (9).

Similar to the history of chemotherapy drug discovery, the search for radiosensitizers has included a large diversity of compounds, ranging from caffeine to nanoparticles and quantum dots (10,11). Notably, chemotherapy has become well-established as having a synergistic effect with radiotherapy, enhancing efficacy (12,13). The academic growth of radiosensitizers is substantial, spanning sensitization of many different radiotherapy modalities and cancer types (14). Given continued progress and diversification of radiosensitizer research, it is valuable to quantify and assess the development of this academic field. Bibliometrics is a data science-based approach that involves analysis of publications to characterize trends within scholarly areas (15). This often involves investigation of authors, journals, institutions and countries and their trends.

## Materials and Methods

### 2.1 Data Collection

A bibliometric analysis was performed for radiosensitizer research from 1956 to 2024. The Web of Science Database was queried for radiosensitizer publications with the following search strategy: ‘radiosensitizer’ OR ‘radiation sensitizer’ OR ‘radio-sensitization’ OR ‘radio-enhancer’ OR ‘radiosensitizing agent’, OR ‘radiosensitization’. Subgroup analysis was then performed by cancer type with additional key words for brain, lung, ear nose and throat (ENT), gastrointestinal (GI), gynecology, genitourinary and breast cancers (Appendix A). Articles from 2025 were initially excluded from the analysis. Conference proceedings and published abstracts were included in the analysis, alongside full-length articles.

### 2.2 Data analysis and Visualization

For analysis of the most cited publications within each organ system and the most cited primary studies in radiosensitization research, reviews, commentaries, guidelines, meta-analyses, and studies which did not investigate the relevant organ systems were excluded. For analysis of the most cited review publications, reviews, commentaries and meta-analyses were included, while all other article types were excluded.

Linear regression was performed to determine the association between year and publication count for global and subgroup trends, with a starting year of 2001. The Bibliometrix package in R was used for various levels of our analysis including elucidating author impacts and trends, productivity of countries, and thematic/key word analysis. VOS Viewer was also used to determine the co-occurrence of keywords, with a threshold value of 5 occurrences. The 10 most co-occurring words in the VOS-Viewer co-occurrences map were removed to visually demonstrate finer trends in keywords/topics without distorting the effects due to the presence of search terms in the keywords list. For the creation of the key word thematic map in Bibliometrix, the top 4,000 key words were utilized, accounting for the presence of synonyms. Generic field terms such as ‘cancer’, ‘cell’, and ‘radiation’ were also excluded from the figure to generate informative thematic plots.

## Results

### 3.1. Global Trend in Radiosensitizer Publications

Our initial search yielded 12,690 records, with publication year ranging from 1956 to 2024. The number of publications per year increased over time, with 1 report in 1956 to 532 reports in 2024. The sharpest increase occurred beginning in 2001, with 215 publications in 2001 (Slope = 17.9, R^2^ =0.93, p<0.001). The trend from 2001 to 2024 is visualized in Figure 1.

**Figure 1:**
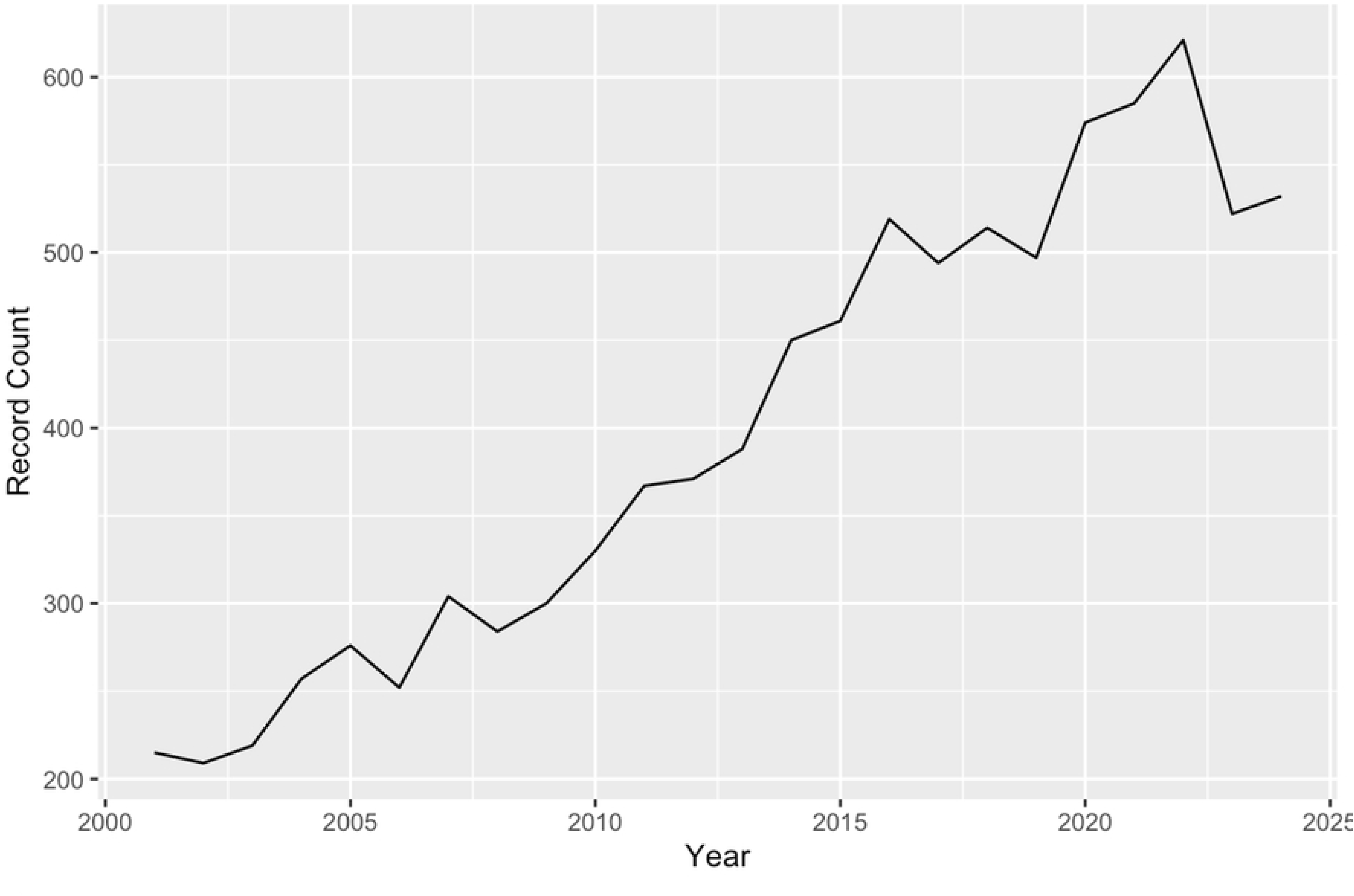
Trend in global radiosensitizer publications over time.

**Figure 2:**
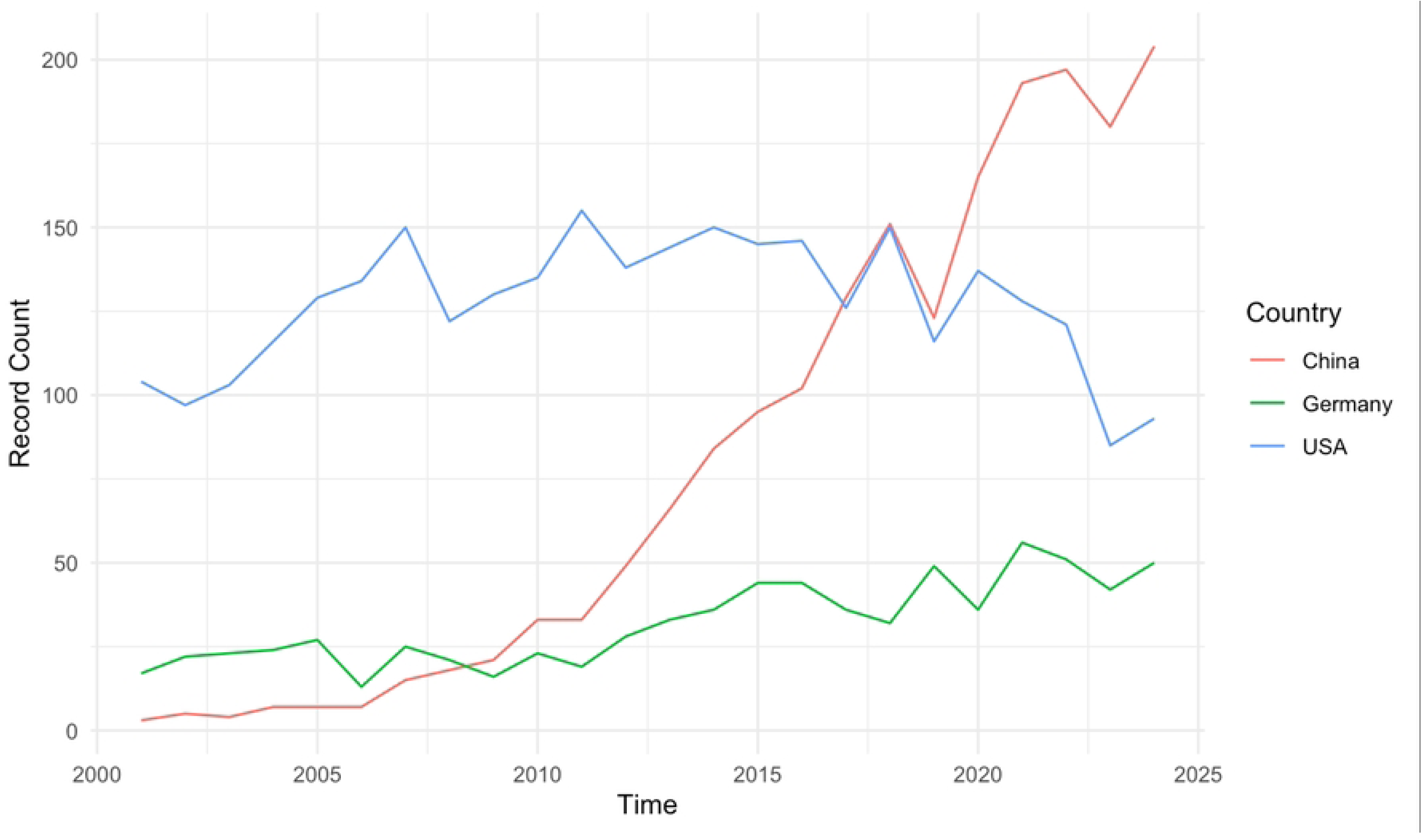
Annual rate of radiosensitizer publications between 2001 and 2024 time for China, Germany, and the United States.

**Figure 3:**
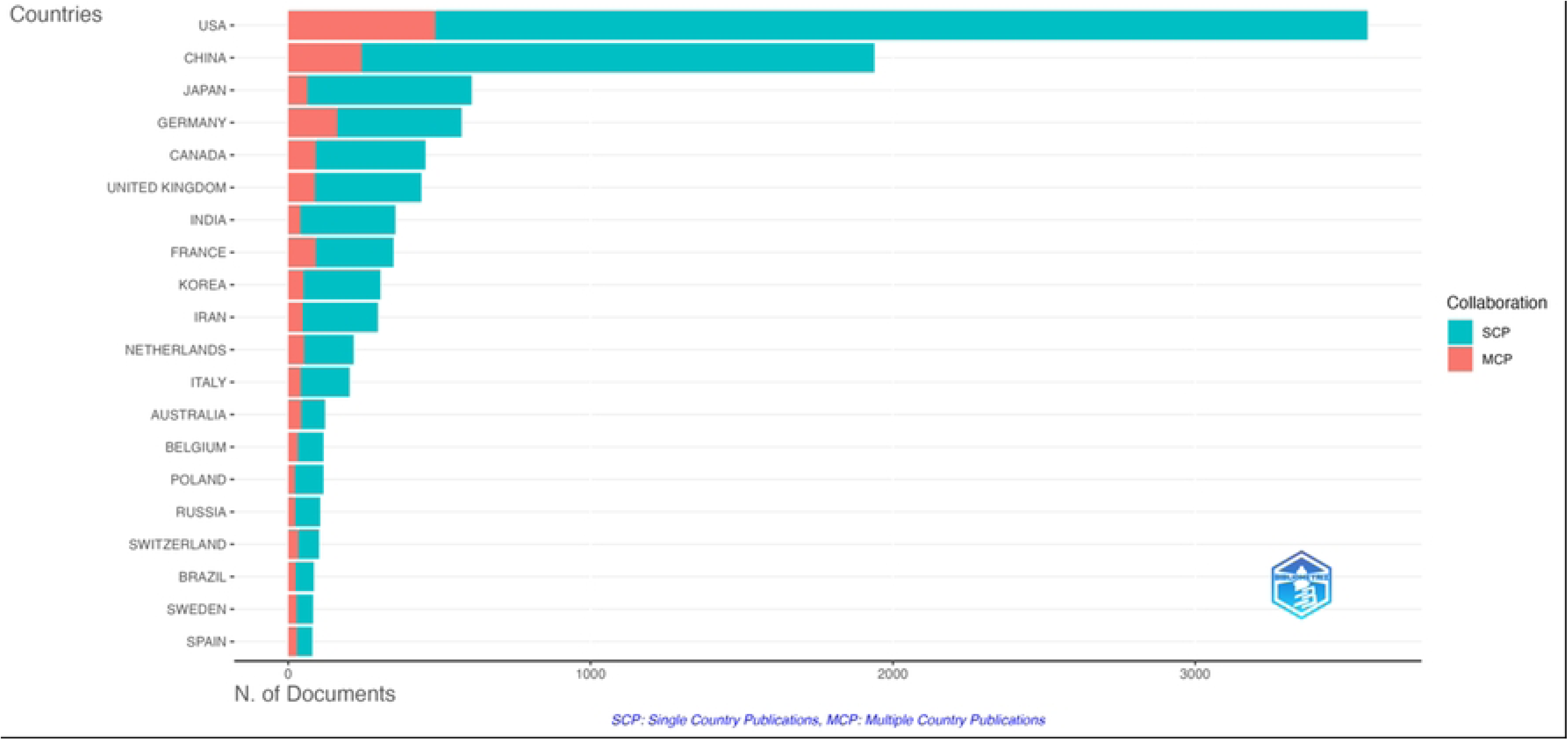
Countries of origin of corresponding authors for radiosensitizer publications. Publications within each country were divided into publications with authors from only one country (SCP) and publications with authors from multiple countries (MCP).

**Figure 4:**
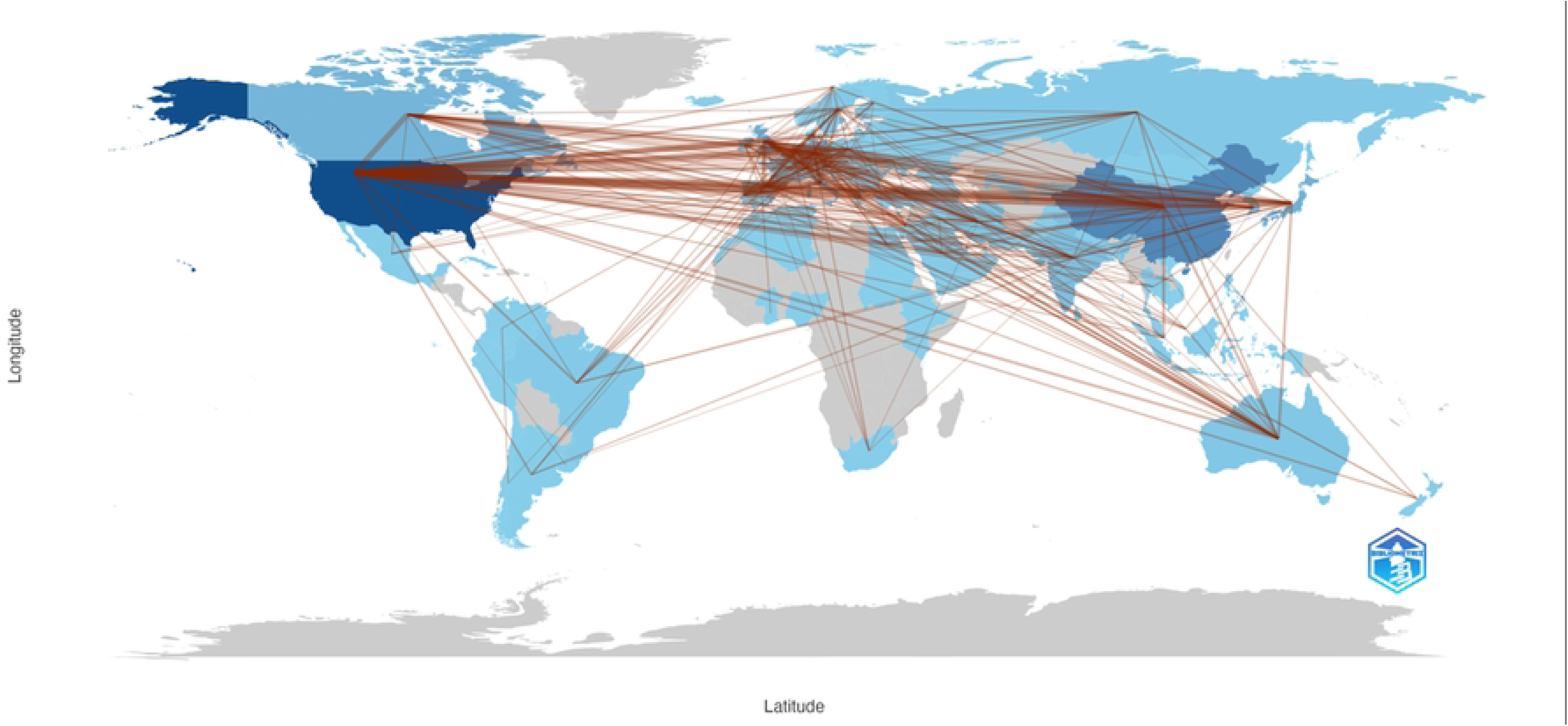
Map of international collaboration through co-authorship for radiosensitizer publications. Countries are color-coded for productivity in terms of record count. Darker shades of blue represent greater productivity, while grey indicates 0 research output from the region.

**Figure 5:**
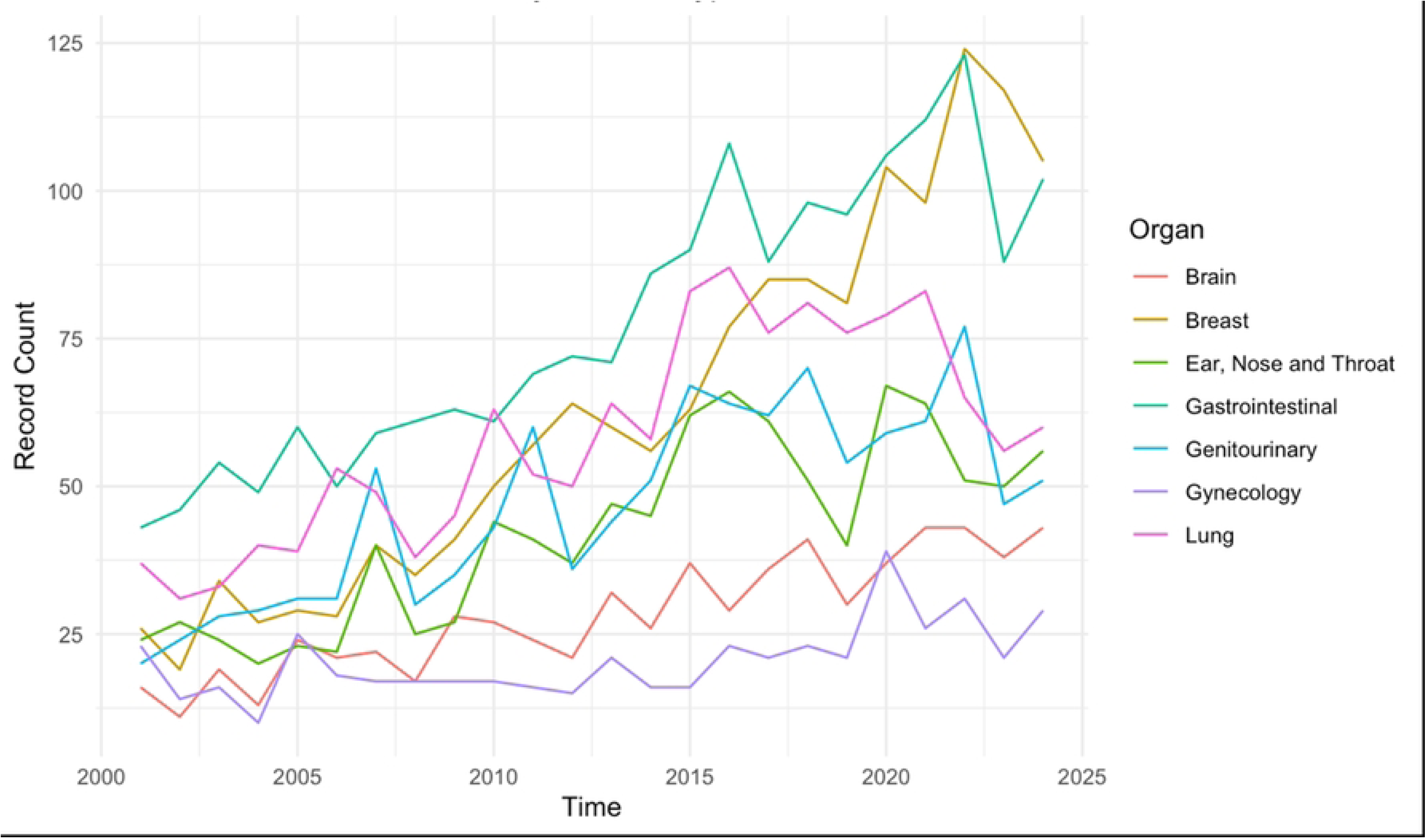
The trend of radiosensitizer publications of each organ type with time.

**Figure 6:**
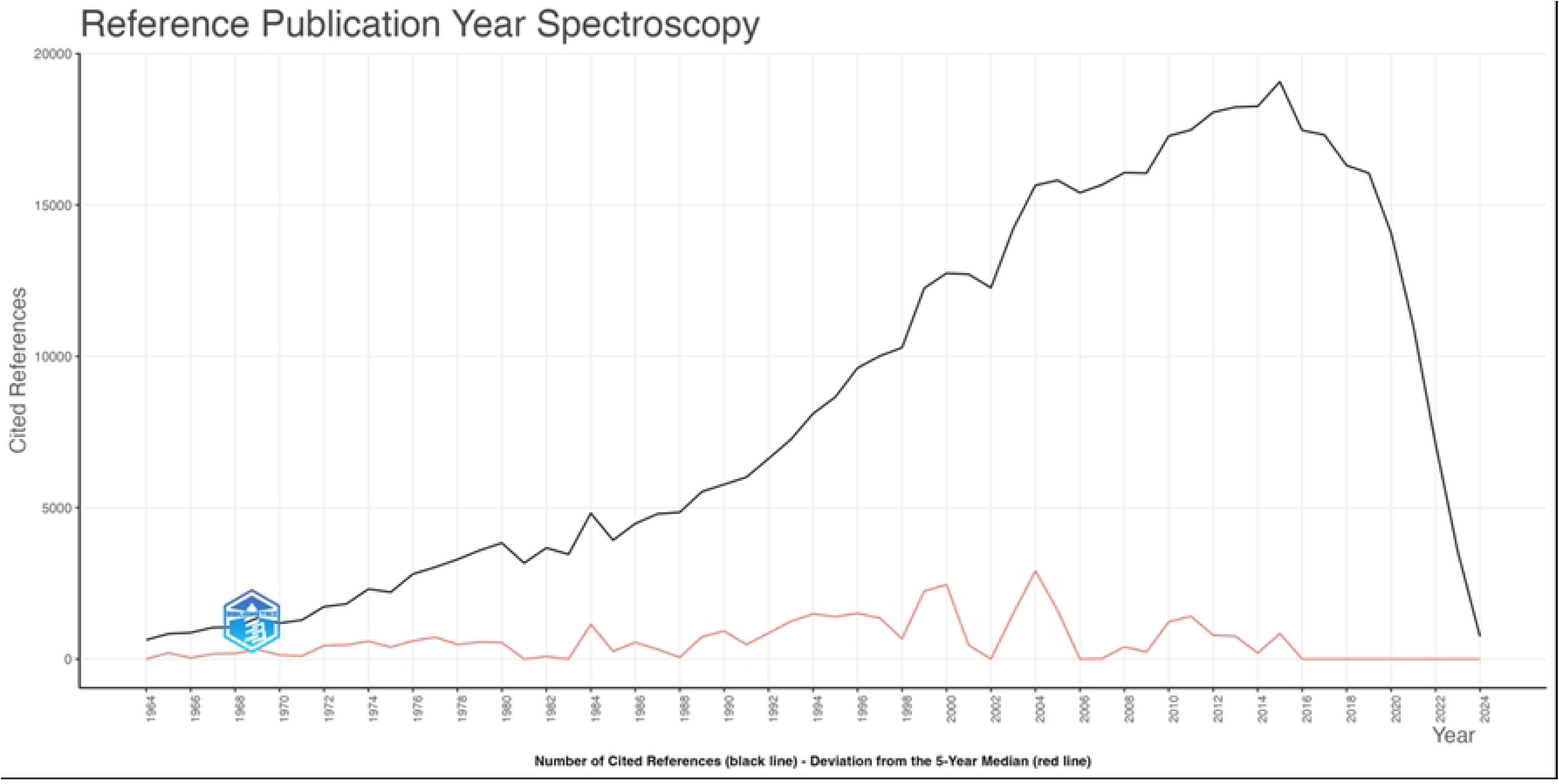
Reference year spectroscopy plot for global radiosensitizer publications.

**Figure 7:**
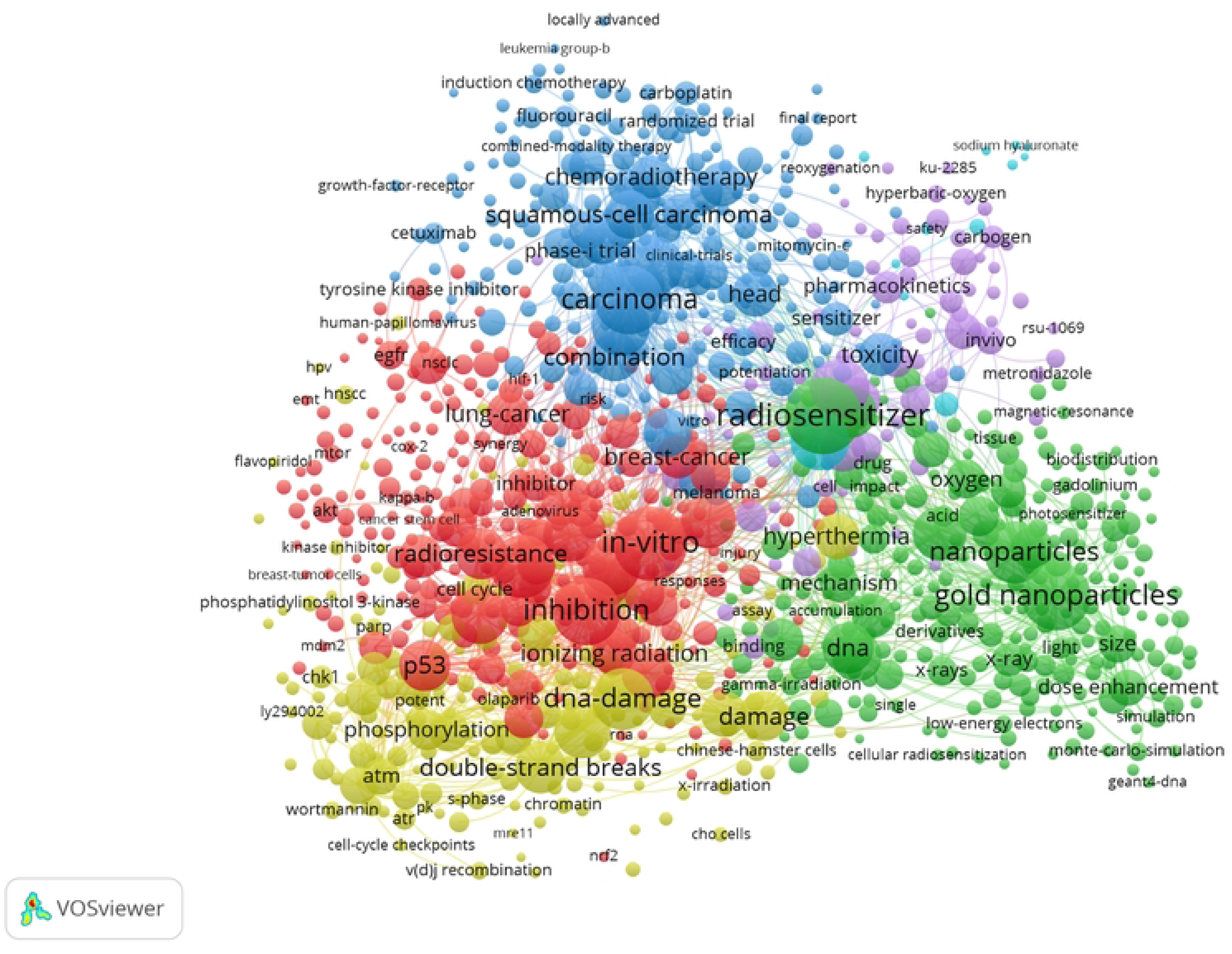
VOS Viewer network visualization of co-occurrences of keywords.

**Figure 8:**
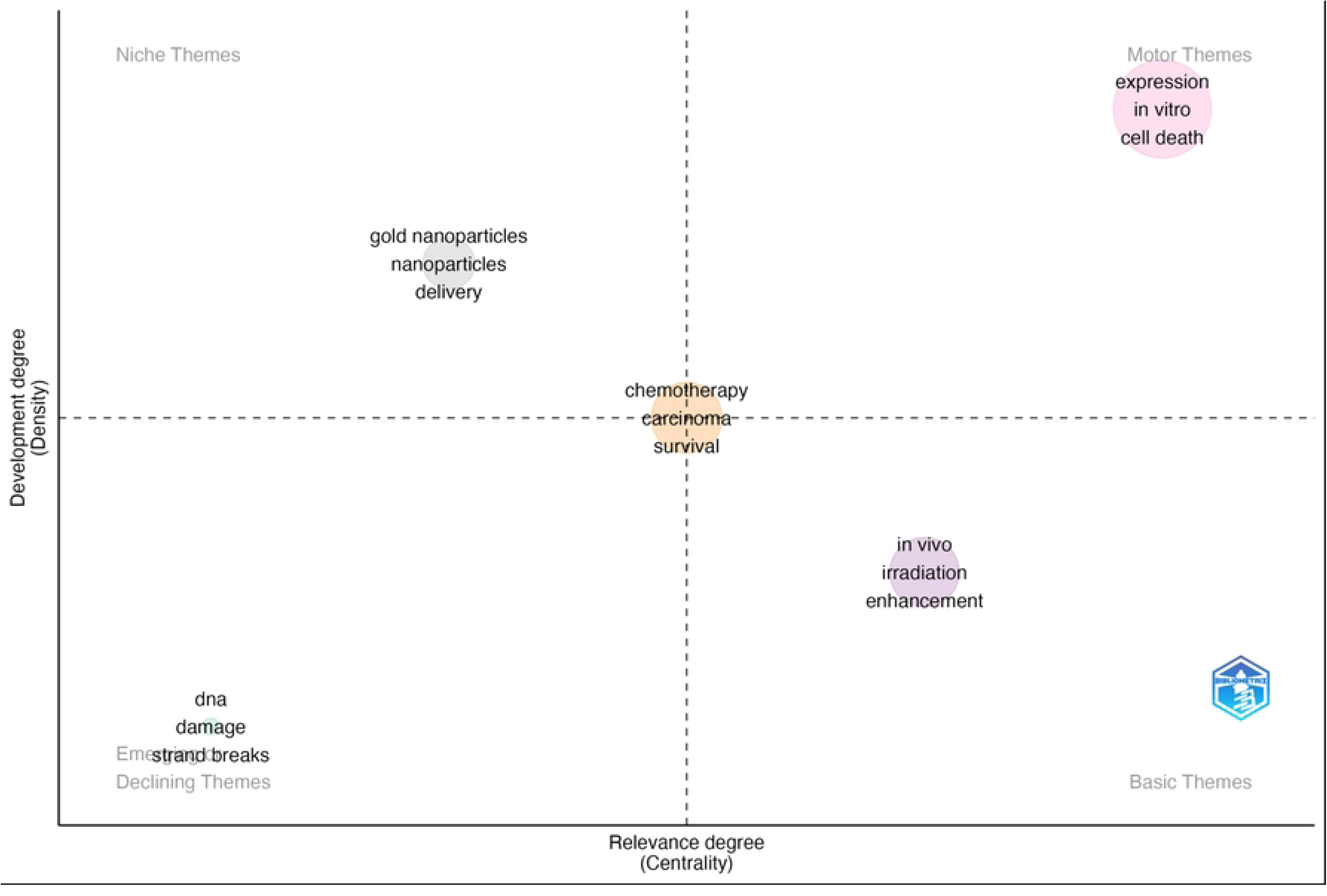
Thematic plot of key topics derived from key word analysis and their dynamics.

2022 was the most productive year for radiosensitizer research, accounting for 4.89% of all included publications. The most highly represented journals were *The International Journal of Radiation Oncology Biology Physics* (n=986, 7.70%), *Radiation Research* (n=483, 3.08%), *Cancer Research* (n=419, 3.30%), *International Journal of Radiation Biology* (n=407, 3.21%), and *Radiotherapy and Oncology* (n=328, 2.585%). The most highly represented affiliated institutions were The University of Texas System (n=459, 3.62%), Harvard University (n=331, 2.61%), The University of California System (n=331, 2.61%), the UTMD Anderson Cancer Center (n=295, 2.33%) and the National Institutes of Health (n=277, 2.18%). The most productive individual authors with regard to total record count were Lawrence, TS (n=129, 1.10%), Wang, Y (n=95, 0.75%), and Kinsella, TJ (n=73, 0.58%).

The ten most highly represented countries were the United States of America (n=4420, 34.83%), China (n=1952, 15.38%), Germany (n=883, 6.96%), Japan (n=861, 6.79%), England (n=845, 6.66%), Canada (n=743, 5.86%), France (n=553, 4.36%), India (n=440, 3.47%), South Korea (n=341, 2.69%), and the Netherlands (n=332, 2.62%). The United States, China and Germany all demonstrated increases in annual publication rate over time, with China showing a significantly higher increase in rate (R^2^ =0.92, p<0.001).

Most corresponding authors were from institutions based in the United States, China, Japan, Germany and Canada, consistent with the representation of these countries in overall publication count. Additionally, the most productive countries in radiosensitizer research in terms of total record count appear to more frequently engage in collaboration with one another. By comparison, less productive countries have fewer collaborations with the most productive countries.

### 3.2. Cancer Type Subgroup Analysis of Trend in Radiosensitizer Publications

#### 3.2.1 Trend in Brain Radiosensitizer Publications

A bibliometric search for brain-related radiosensitizer publications yielded 837 results. Brain-related radiosensitizer publications increase with time (Slope = 1.2696, R^2^ = 0.8272, p<0.001). The most highly represented journals were *The International Journal of Radiation Oncology Biology Physics* (n=73, 8.72%), *Journal of Neuro-Oncology* (n=31, 3.704%), *Neuro-Oncology* (n=22, 2.63%), *Cancer Research* (n=18, 2.15%), and *Clinical Cancer Research* (n=18, 2.15%). The most highly represented countries were the United States (n=432, 51.61%), China (n=98, 11.71%), France (n=59, 7.05%), Germany (n=51, 6.09%), and England (n=44, 5.26%). The most productive authors in terms of total record count were Mehta MP (n=12, 1.43%), Lux F (n=11, 1.31%), and Camphausen K (n=10, 1.20%).

#### 3.2.2 Trend in Lung Radiosensitizer Publications

A bibliometric search for lung-related radiosensitizer publications yielded 1,696 results. Lung radiosensitizer publications increase with time (Slope = 1.198, R^2^ = 0.5794, p<0.001). The most highly represented journals were *Anticancer Research* (n=28, 1.65%), *British Journal of Cancer* (n=21, 1.24%), *Cancer Letters* (n=24, 1.42%), *Cancer Research* (n=46, 2.71%), and *Clinical Cancer Research* (n=38, 2.24%). The most highly represented countries were The United States (n=666, 39.27%), China (n=348, 20.51%), Germany (n=133, 7.84%), Japan (n=102, 6.01%), and South Korea (n=89, 5.25%). The most productive authors in terms of total record count were Choy H (n=39, 2.30%), Wang Y (n=27, 1.59%), and Lu B (n=22, 1.30%).

#### 3.2.3 Trend in ENT Radiosensitizer Publications

A bibliometric analysis of ENT-related radiosensitizer publications yielded 1, 213 results. ENT radiosensitizer publications increase with time (Slope = 1.8278, R^2^ = 0.6756, p <0.001). The most highly represented journals were *The International Journal of Radiation Oncology Biology Physics* (n=111, 9.15%), *Radiotherapy and Oncology* (n=74, 6.101%), *Clinical Cancer Research* (n=31, 2.56%), *Cancer Research* (n=30, 2.47%) and *Cancers* (n=24, 1.98%). The most highly represented countries were The United States (n=421, 34.70%), China (n=206, 16.98%), Germany (n=124, 10.22%), Japan (n=78, 6.43%), England (n=77, 6.35%). The most prolific authors in terms of total record count were Overgaard J (n=25, 2.06%), Cordes N (n=19, 1.57%), and Lawrence TS (n=18, 1.48%).

#### 3.2.4 Trend in GI Radiosensitizer Publications

A bibliometric analysis of GI-related radiosensitizer publications yielded 2,174 results. GI publications increase with time (Slope = 3.0830, R^2^ = 0.8569, p<0.001). The most highly represented journals were the *International Journal of Radiation Oncology Biology Physics* (n=201, 9.25%), *Cancer Research* (n=77, 3.54%), *Clinical Cancer Research* (n=60, 2.76%), *Radiotherapy and Oncology* (n=55, 2.53%), and *The International Journal of Radiation Biology* (n=37, 1.70%). The most highly represented countries were The United States (n=820, 37.72%), China (n=440, 20.24%), Japan (n=178, 8.19%), Germany (n=156, 7.18%), and South Korea (n=87, 4.00%). The most prolific authors in terms of total record count were Lawrence TS (n=87, 4.00%), Kinsella TJ (n=36, 1.66%), and Davis MA (n=26, 1.20%).

#### 3.2.5 Trend in Gynecological Radiosensitizer Publications

A bibliometric search for gynecological-related radiosensitizer publications yielded 667 publications. Gynecological publications increase with time (Slope = 0.5374, R^2^ = 0.334, P<0.01). The most highly represented journals were *The International Journal of Radiation Oncology Biology Physics* (n=54, 8.10%), *Gynecologic Oncology* (n=45, 6.74%), *Cancer Research* (n=17, 2.55%), *Radiotherapy and Oncology* (n=16, 2.34%), and *International Journal of Gynecological Cancer* (n=15, 2.25%). The most represented countries were the United States (n=256, 38.38%), China (n=125, 18.74%), Japan (n=45, 6.75%), Canada (n=41, 6.15%) and Germany (n=33, 4.95%). The most prolific authors in terms of total record count were Chen TF (n=16, 2.34%), Franken NAP (n=13, 1.95%), and Crezee J (n=11, 1.65%).

#### 3.2.6 Trend in Breast Cancer Radiosensitizer Publications

A bibliometric search for breast cancer related radiosensitizer publications yielded 1,170 results. Breast cancer radiosensitizer publications increase with time (Slope = 4.2874, R^2^ = 0.9291, p<0.001). The most represented journals were *The International Journal of Radiation Oncology Biology Physics* (n=111, 6.49%), *Cancer Research* (n=62, 3.63%), *Radiation Research* (n=42, 2.47%), *Radiotherapy and Oncology* (n=37, 2.16%), and *Cancers* (n=33, 1.93%). The most represented countries were The United States (n=612, 35.79%), China (n=314, 18.36%), Germany (n=125, 7.31%), Iran (n=107, 6.26%), and Japan (n=102, 5.97%). The most prolific authors in terms of total record count were Pierce LJ (n=32, 1.87%), Wilder-Romans K (n=26, 1.52%), Ogawa Y (n=25, 1.46%), Speers C (n=25, 1.46%), Liu ML (n=23, 1.35%).

#### 3.2.7 Composite Trend Analysis

Publications related to radiosensitizers in the context of breast cancer have increased most quickly with time compared to all other cancer types, while radiosensitizer publications in the context of gynecological cancers have seen the slowest growth (p<0.0001).

### 3.3. Cancer Type Subgroup Analysis of the Top Manuscripts

#### 3.3.1 The Most Cited Manuscripts for Brain Radiosensitizers

The most highly cited study for brain cancer related radiosensitizer publications was “Taxol – A Novel Radiation Sensitizer” published by Tishler, RB and Hall, EJ in the *International Journal of Radiation Oncology Biology Physics* in 1992 with 319 citations to date. The radiosensitizers investigated in the top 10 brain-related publications include Taxol, ATM kinase inhibitors, Poly(ADP-ribose) polymerase (PARPis), valproic acid, gold nanoparticles, gadolinium nanoparticles, cisplatin, and mTOR inhibitors. Studies were primarily published in oncology or radiation specific journals, with 3 studies as exceptions, published in *ACS Nano*, *Science Advances*, and *PLOS One*. 7 studies utilized xenograft glioma models, while 3 publications utilized syngeneic murine glioma models. The oldest publication was a clinical trial for cisplatin as a radiosensitizer for malignant brain tumors, published in 1982 (Table 1).

**Table 1:**
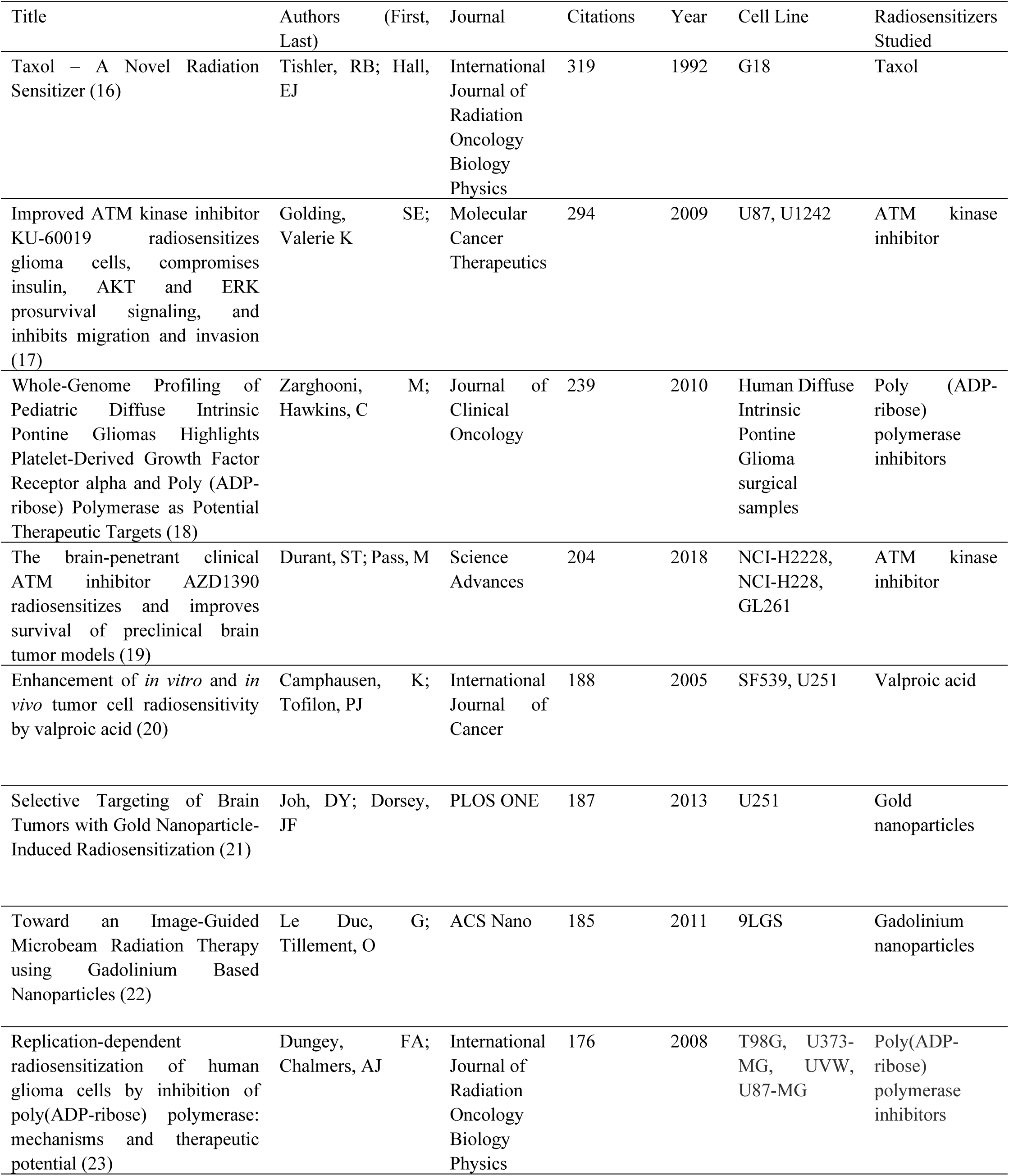
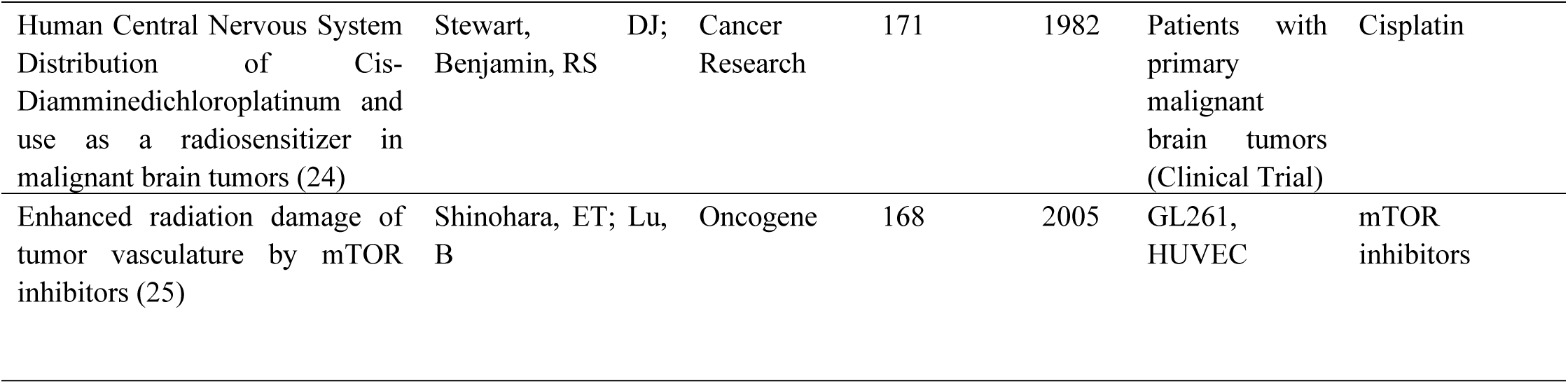
The 10 Most Cited Brain Cancer Focused Radiosensitizer Publications.

#### 3.1.2 The Most Cited Manuscripts for Lung Cancer Radiosensitizers

The most highly cited study for lung cancer related radiosensitizer publications was “Inhibition of ATM and ATR kinase activities by the radiosensitizing agent, caffeine”, published by Sarkaria JN and Abraham, RT in *Cancer Research* in 1999 with 946 citations to date. The radiosensitizers discussed in the 5 most cited lung publications include caffeine (as an ATM/ATR inhibitor), wortmannin, cisplatin, Taxol and PARPis. All 5 studies were published in cancer-specific journals. 3 out of the 5 studies were published prior to the 21st century, with the oldest publication studying Taxol in 1994 (Table 2).

**Table 2:** The 5 Most Cited Lung Cancer Focused Radiosensitizer Publications.

| Title | Authors (First, Last) | Journal | Citations | Year | Radiosensitizers Studied |
| --- | --- | --- | --- | --- | --- |
| Inhibition of ATM and ATR kinase activities by the radiosensitizing agent, caffeine (26) | Sarkaria, JN; Abraham, RT | Cancer Research | 946 | 1999 | Caffeine |
| Inhibition of phosphoinositide-3-kinase related kinases by the radiosensitizing agent wortmannin (27) | Sarkaria, JN; Abraham, RT | Cancer Research | 501 | 1998 | Wortmannin |
| Randomized phase II study of cisplatin with gemcitabine or paclitaxel or vinorelbine as induction chemotherapy followed by concomitant chemoradiotherapy for stage IIB non-small-cell lung cancer: Cancer and leukemia group B study 9431(28) | Vokes, EE; Green, MR | Journal of Clinical Oncology | 290 | 2002 | Cisplatin |
| In-vitro studies of Taxol as a radiation sensitizer in human tumor cells (29) | Liebmann, J; Mitchell, JB | Journal of the National Cancer Institute | 278 | 1994 | Taxol |
| Inhibition of poly (ADP-ribose) polymerase enhances cell death and improves tumor growth delay in irradiated lung cancer models (30) | Albert, JM; Lu, B | Clinical Cancer Research | 239 | 2007 | Poly(ADP-ribose) polymerase inhibitors |

#### 3.1.2. The Most Cited Manuscripts for ENT Cancer Radiosensitizers

The most highly cited study for ear, nose and throat cancer related radiosensitizer publications was “Association of reactive oxygen species and radio resistance in cancer stem cells”, published by Diehn M and Clarke MF in *Nature* in 2009 with 1,988 citations to date. The 5 most cited ENT studies examined inhibition of ROS scavengers in cancer stem cells with buthionine sulfoximine, nimorazole, EGFR inhibitors, cisplatin, and beta - 1 Integrin inhibitors.

The 5 most cited studies were published in cancer or radiation specific journals with two exceptions – one in *Nature*, and one in the *Journal of Clinical Investigation*. The oldest study was a clinical trial of nimorazole as a radiosensitizer for head and neck cancer patients, published in 1998 (Table 3).

**Table 3:** The 5 Most Cited ENT Cancer Focused Radiosensitizer Publications.

| Title | Authors(First;Last) | Journal | Citations | Year | Radiosensitizers Studied |
| --- | --- | --- | --- | --- | --- |
| Association of reactive oxygen species and radioresistance in cancer stem cells (31) | Diehn, M; Clarke, MF | Nature | 1,988 | 2009 | Inhibition of ROS scavengers in cancer stem cells |
| A randomized double-blind phase III study of nimorazole as a hypoxic radiosensitizer of primary radiotherapy in supraglottic larynx and pharynx carcinoma, results of the Danish Head and Neck Cancer Study (DAHANCA) protocol 5-85(32) | Overgaard, J; Jørgensen, K | Radiotherapy and Oncology | 464 | 1998 | Nimorazole |
| Local recurrence in head and neck cancer: relationship to radiation resistance and signal transduction (33) | Gupta, AK; Muschel, RJ | Clinical Cancer Research | 312 | 2002 | EGFR inhibitors |
| Combined postoperative radiotherapy and weekly cisplatin infusion for locally advanced head and neck carcinoma: Final report of a randomized trial (34) | Bachaud, JM; Daley-Schvartz, N | International Journal of Radiation Oncology Biology Physics | 264 | 1996 | Cisplatin |
| β1 Integrin/FAK/cortactin signaling is essential for human head and neck cancer resistance to radiotherapy (35) | Eke, I; Cordes, N | Journal of Clinical Investigation | 183 | 2012 | β1 Integrin inhibitors |

#### 3.1.3 The Most Cited Manuscripts for GI Cancer Radiosensitizers

The most cited study for gastrointestinal cancer related radiosensitizer publications was “Pharmacological requirements for obtaining sensitization of human tumor cells in vitro to 5-fluorouracil or ftorafur and x rays”, published by Byfield, JE and Kulhanian, F in the *International Journal of Radiation Oncology Biology Physics* in 1982 with 354 citations to date. The 5 most cited gastrointestinal studies examined radiosensitizers of 5-fluorouracil, DNA dependent protein kinase inhibitors, NF-kB inhibitors, Taxol, and diflourodeoxycytidine. The 5 most cited studies were all published in radiation- or oncology-specific journals. Three out of 5 studies were published in the 20th century, with the oldest study, published in 1982, being the most cited (Table 4).

**Table 4:** The 5 Most Cited GI Cancer Focused Radiosensitizer Publications.

| Title | Authors<br>(First; Last) | Journal | Citations | Year | Radiosensitizers<br>Studied |
| --- | --- | --- | --- | --- | --- |
| Pharmacologic requirements for obtaining sensitization of human tumor cells in vitro to combined 5-fluorouracil or ftorafur and x rays (36) | Byfield, JE;<br>Kulhanian,<br>F | International Journal of Radiation<br>Oncology Biology Physics | 354 | 1982 | 5-fluorouracil, ftorafur |
| Preclinical evaluation of a potent novel DNA dependent protein kinase inhibitor NU7441 (37) | Zhao Y;<br>Curtin, NJ | Cancer Research | 350 | 2006 | DNA-dependent<br>protein kinase<br>inhibitors |
| Enhancement of radiosensitivity by proteasome inhibition: Implications for a role of NF-kB (38) | Russo, SM;<br>Cusack, JC | International Journal of Radiation<br>Oncology Biology Physics | 291 | 2001 | NF-kB inhibitors |
| In-vitro studies of Taxol as a radiation sensitizer in human tumor cells (29) | Liebmann,<br>J; Mitchell,<br>JB | Journal of the National Cancer<br>Institute | 278 | 1994 | Taxol |
| Radiosensitization of pancreatic cancer cells by 2',2'-difluoro-2'deoxyctidine (39) | Lawrence,<br>TS;<br>Shewach,<br>DS | International Journal of Radiation<br>Oncology Biology Physics | 298 | 1996 | Diflourodeoxycytidine |

#### 3.1.4 The Most Cited Manuscripts for Gynecological Cancer Radiosensitizers

The most cited study for gynecologic cancer related radiosensitizer publications was “Pharmacological requirements for obtaining sensitization of human tumor cells in vitro to 5-fluorouracil or ftorafur and x rays”, by Byfield, JE and Kulhanian, F. It was published in the *International Journal of Radiation Oncology Biology Physics* in 1982 with 354 citations to-date.

The 5 most cited gynecologic cancer related studies investigated radiosensitization approaches involving the use of 5-fluorouracil, Ad5-CD/TKrep-mediated double suicide gene therapy, geldanamycin and its analogs, thio-glucose bound gold nanoparticles, and curcumin. Publications were split between clinical radiation/oncology-specific journals, and basic science journals such as *Human Gene Therapy*, *Nanotechnology* and *Molecular Pharmacology*. Besides the most cited study, the other 4 studies comprising the top 5 were published in the 21st century (Table 5).

**Table 5:** The 5 Most Cited Gynecological Cancer Focused Radiosensitizer Publications.

| Title | Authors (First; Last) | Journal | Citations | Year | Radiosensitizers Studied |
| --- | --- | --- | --- | --- | --- |
| Pharmacologic requirements for obtaining sensitization of human tumor cells in vitro to combined 5-fluorouracil or ftorafur and x rays (36) | Byfield, JE; Kulhanian, F | International Journal of Radiation Oncology Biology Physics | 354 | 1982 | 5-fluorouracil, ftorafur |
| Double suicide gene therapy augments the antitumor activity of a replication competent lytic adenovirus through enhanced cytotoxicity and radiosensitization (40) | Rogulski, KR; Freytag, SO | Human Gene Therapy | 155 | 2000 | Ad5-CD/TKrep-mediated double suicide gene therapy |
| Geldanamycin and 17-allylamino-17-demethoxygeldanamycin potentiate the in vitro and in vivo radiation response of cervical tumor cells via the heat shock protein 90-mediated intracellular signaling and cytotoxicity (41) | Bisht, KS; Gius, D | Cancer Research | 149 | 2003 | Geldanamycin, Geldanamycin analogs |
| Thio-glucose bound gold nanoparticles enhance radio-cytotoxic targeting of ovarian cancer (42) | Geng, F; Kong, BH | Nanotechnology | 145 | 2011 | Thio-glucose bound gold nanoparticles |
| The chemo preventative agent curcumin is a potent radiosensitizer of human cervical tumor cells via increased reactive oxygen species production and overactivation of the mitogen activated protein kinase pathway (43) | Javvadi, P; Koumenis, C | Molecular Pharmacology | 132 | 2008 | Curcumin |

#### 3.1.5 The Most Cited Manuscripts for Genitourinary Cancer Radiosensitizers

The most cited study for genitourinary cancer related radiosensitizer publications was “Cell-specific radiosensitization by gold nanoparticles at megavoltage radiation energies”, published by Jain, S and Hirst, DG in the *International Journal of Radiation Oncology Biology Physics* in 2011 with 371 citations to date. The radiosensitizers studied in the 5 most cited publications were gold nanoparticles, paclitaxel, mTOR inhibitors, and curcumin. Most studies were published in oncology or radiation specific journals with the exception of two studies, published in *Cell Death and Disease* and *Nanotechnology* (Table 6).

**Table 6:** The 5 Most Cited Genitourinary Cancer Focused Radiosensitizer Publications.

| Title | Authors (First; Last) | Journal | Citations | Years | Radiosensitizers Studied |
| --- | --- | --- | --- | --- | --- |
| Cell-specific radiosensitization by gold nanoparticles at megavoltage radiation energies (44) | Jain, S; Hirst, DG | International Journal of Radiation Oncology Biology Physics | 371 | 2011 | Gold nanoparticles |
| Significant Activity of Paclitaxel in Advanced Transitional Cell Carcinoma of the Urothelium – A Phase II Trial of the Eastern Cooperative Oncology Group (45) | Roth, BJ; Loehrer, PJ | Journal of Clinical Oncology | 290 | 1994 | Paclitaxel (Taxol) |
| Gold nanoparticle sensitize radiotherapy of prostate cancer cells by regulation of the cell cycle (46) | Roa, W; Xing, JZ | Nanotechnology | 264 | 2009 | Gold nanoparticles |
| PI3K/Akt/mTOR pathway inhibitor enhance radiosensitivity in radioresistant prostate cancer cells through including apoptosis, reducing autophagy, suppressing NHEJ and HR repair pathways (47) | Chang, L; Li, Y | Cell Death and Disease | 255 | 2014 | mTOR inhibitors |
| Curcumin, a dietary component, has anticancer, chemosensitization, and radiosensitization effects by down regulation the MDM2 oncogene through the PI3K/mTOR/ETS2 pathway (48) | Li, M; Zhang, RW | Cancer Research | 250 | 2007 | Curcumin |

#### 3.3.6 The Most Cited Manuscripts for Breast Cancer Radiosensitizers

The most cited study for breast cancer related radiosensitizer publications was “Association of reactive oxygen species and radioresistance in cancer stem cells”, published by Diehn, M and Clarke, MF in *Nature* in 2009 with 1,988 citations to date. The radiosensitizers studied in the 5 most cited publications were inhibition of ROS scavengers in cancer stem cells with buthionine sulfoximine, gold nanoparticles, Taxol, and hyperthermia treatment. Three out 5 studies were published in oncology or radiology journals, with the 2 studies published in *Nature* and *The Journal of Pharmacy and Pharmacology*. The earliest publication was in 1980 and examined hyperthermia as a radiosensitizing therapy (Table 7).

| Title | Authors (First; Last) | Journal | Citations | Years | Radiosensitizers Studied |
| --- | --- | --- | --- | --- | --- |
| Association of reactive oxygen species and radioresistance in cancer stem cells (31) | Diehn, M; Clarke, MF | Nature | 1,988 | 2009 | Inhibition of ROS scavengers in cancer stem cells |
| Radiotherapy enhancement with gold nanoparticles (49) | Hainfeld JF; Smilowitz, HM | Journal of Pharmacy and Pharmacology | 551 | 2008 | Gold nanoparticles |
| Cell Specific Radiosensitization by Gold Nanoparticles at Megavoltage Radiation Energies (44) | Jain, S; Hirst, DG | International Journal of Radiation Oncology Biology Physics | 372 | 2011 | Gold nanoparticles |
| Simultaneous and Sequential Hyperthermia and Radiation Treatment of an Experimental Tumor and its surrounding normal tissue invivo (50) | Overgaard, J | International Journal of Radiation Oncology Biology Physics | 310 | 1980 | Hyperthermia |
| In-vitro studies of Taxol as a radiation sensitizer in human tumor cells (29) | Liebmann, J; Mitchell, JB | Journal of the National Cancer Institute | 278 | 1994 | Taxol |

### 3.4. Global Citation Analysis

#### 3.4.1 The 20 Most Cited Global Reviews and Basic Investigations for Radiosensitizers

The most cited non-review investigation was “Association of reactive oxygen species and radio resistance in cancer stem cells”, published by Diehn, M and Clarke, MF in *Nature* in 2009 with 1,988 citations to date (Table 8).

**Table 8:** The 20 Most Cited Non-Review Radiosensitizer Publications.

| Title | Authors (First; Last) | Journal | Citations | Year |
| --- | --- | --- | --- | --- |
| Association of reactive oxygen species and radioresistance in cancer stem cells (31) | Diehn, M; Clarke, FM | Nature | 1989 | 2009 |
| Inhibition of ATM and ATR Kinase Activities by the Radiosensitizing Agent, Caffeine (26) | Sarkaria, JN; Abraham, RT | Cancer Research | 946 | 1999 |
| Improved Survival with Bevacizumab in Advanced Cervical Cancer (51) | Tewari, KS; Monk, BJ | The New England Journal of Medicine | 924 | 2014 |
| ATR-mediated checkpoint pathways regulate phosphorylation and activation of human Chk1 (52) | Zhao, H; Piwnicka-Worms, H | Molecular and Cellular Biology | 879 | 2001 |
| P53 Binding protein 1 (53BP1) is an early participant in the cellular response to DNA double strand breaks (53) | Schultz, LB; Halazonetis, TD | Journal of Cell Biology | 732 | 2000 |
| Genetics of human cell lines – incorporation of 5-bromodeoxyuridine and 5-iododeoxyuridine into the deoxyribonucleic acid of human cells and its effect on radiation sensitivity (54) | Djordjevic, B; Szybalski, W | Journal of Experimental Medicine | 641 | 1960 |
| Radiotherapy enhancement with gold nanoparticles (49) | Hainfeld, JF; Smilowitz, HM | Journal of Pharmacy and Pharmacology | 551 | 2008 |
| Glucose-Responsive Sequential Generation of Hydrogen Peroxide and Nitric Oxide for Synergistic Cancer Starving-Like/Gas Therapy (55) | Fan, WP; Chen, XY | Angewandte Chemie-International Edition | 541 | 2017 |
| The cell-cycle checkpoint kinase Chk1 is required for mammalian homologous recombination repair (56) | Sorensen, CS; Helleday, T | Nature Cell Biology | 533 | 2005 |
| Gold Nanoparticles as Radiation Sensitizers in Cancer Therapy (9) | Chithrani, DB; Jaffray, DA | Radiation Research | 532 | 2010 |
| Inhibition of phosphoinositide 3-kinase related kinases by the radiosensitizing agent wortmannin (27) | Sarkaria JN; Abraham, RT | Cancer Research | 501 | 1998 |
| Photoredox catalysis using infrared light via triplet fusion upconversion (57) | Ravetz, BD; Campos, LM | Nature | 481 | 2019 |
| P53-deficient cells rely on ATM and ATR mediated checkpoint signaling through the p38MAPK/MK2 pathway for survival after DNA damage (58) | Reinhardt, HC; Yaffe, MB | Cancer Cell | 478 | 2007 |
| A randomized double-blind phase III study of nimorazole as a hypoxic radiosensitizer of primary radiotherapy in supraglottic larynx and pharynx carcinoma, results of the Danish Head and Neck Cancer Study (DAHANCA) protocol 5-85 (32) | Overgaard, J; Jorgensen, J | Radiotherapy and Oncology | 464 | 1998 |
| Blocked autophagy sensitizes resistant carcinoma cells to radiation therapy (59) | Apel, A; Mayer, A | Cancer Research | 449 | 2008 |
| Polo-like kinase (Plk) 1 depletion induces apoptosis in cancer cells (60) | Liu, XQ; Erikson, RL | Proceedings of the National Academy of Sciences | 439 | 2003 |
| Anticancer chemosensitization and radiosensitization by the novel poly(ADP-ribose) polymerase-1 inhibitor AG14361 (61) | Calabrese, CR; Curtin, NJ | Journal of National Cancer Institute | 425 | 2004 |
| Size-dependent radiosensitization of PEG coated gold nanoparticles for cancer radiation therapy (62) | Zhang, XD; Liang, XJ | Biomaterials | 423 | 2012 |
| Electron-Affinic Sensitization – VII. A correlation between structures, one-electron reduction potentials, and efficiencies of nitroimidazoles as hypoxic cell radiosensitizers (63) | Adams, GE; Watts, ME | Radiation Research | 403 | 1976 |
| Nanoscale metal organic frameworks for combined photodynamic and radiation therapy in cancer treatment (64) | Liu, JJ; Liu, Z | Biomaterials | 388 | 2016 |

The most cited review article was “Molecular Photovoltaics” published by Hagfeldt, A and Gratzel, M *in Accounts of Chemical Research* in 2009, with 2,613 citations to date (Table 9).

**Table 9:**
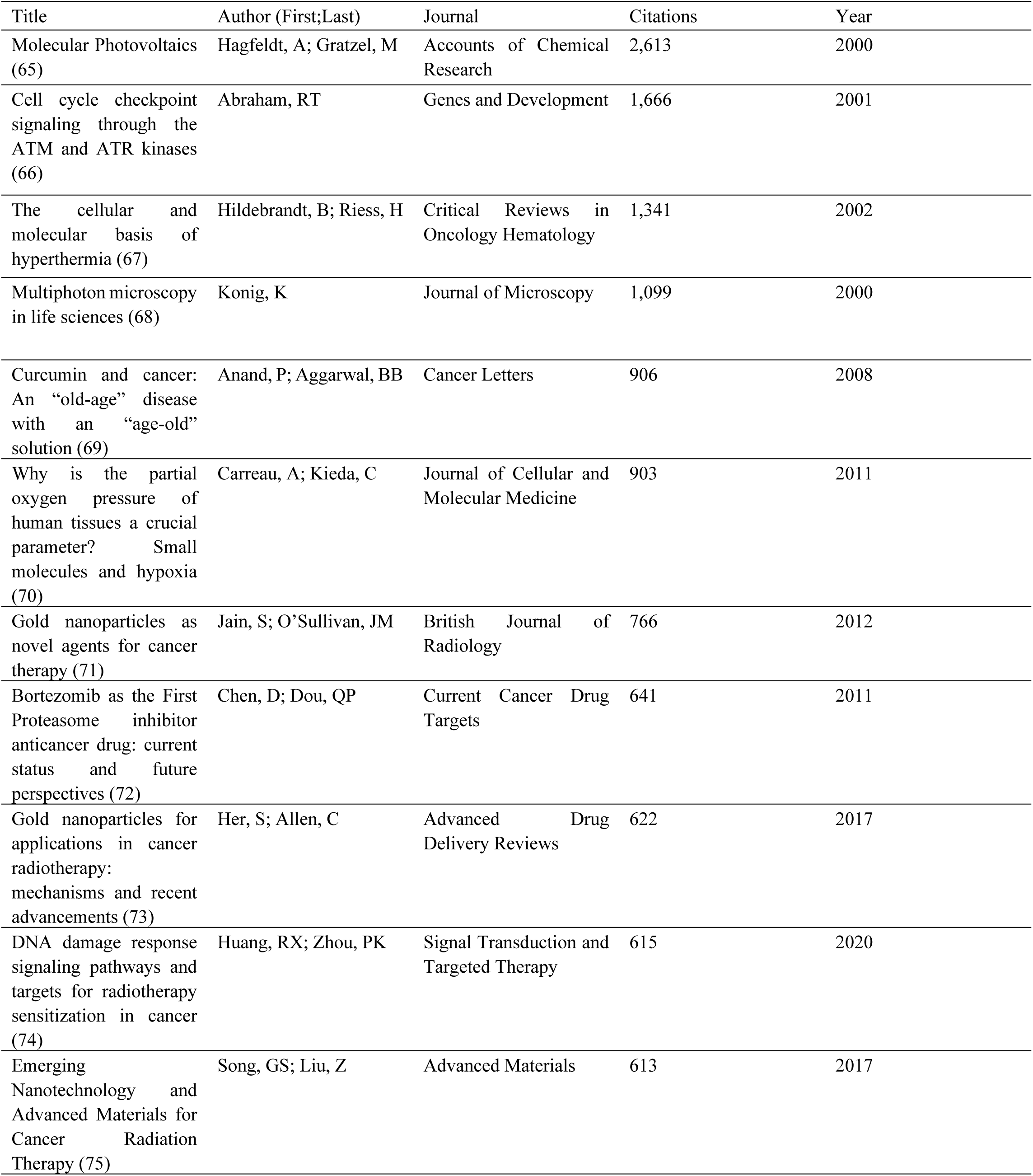
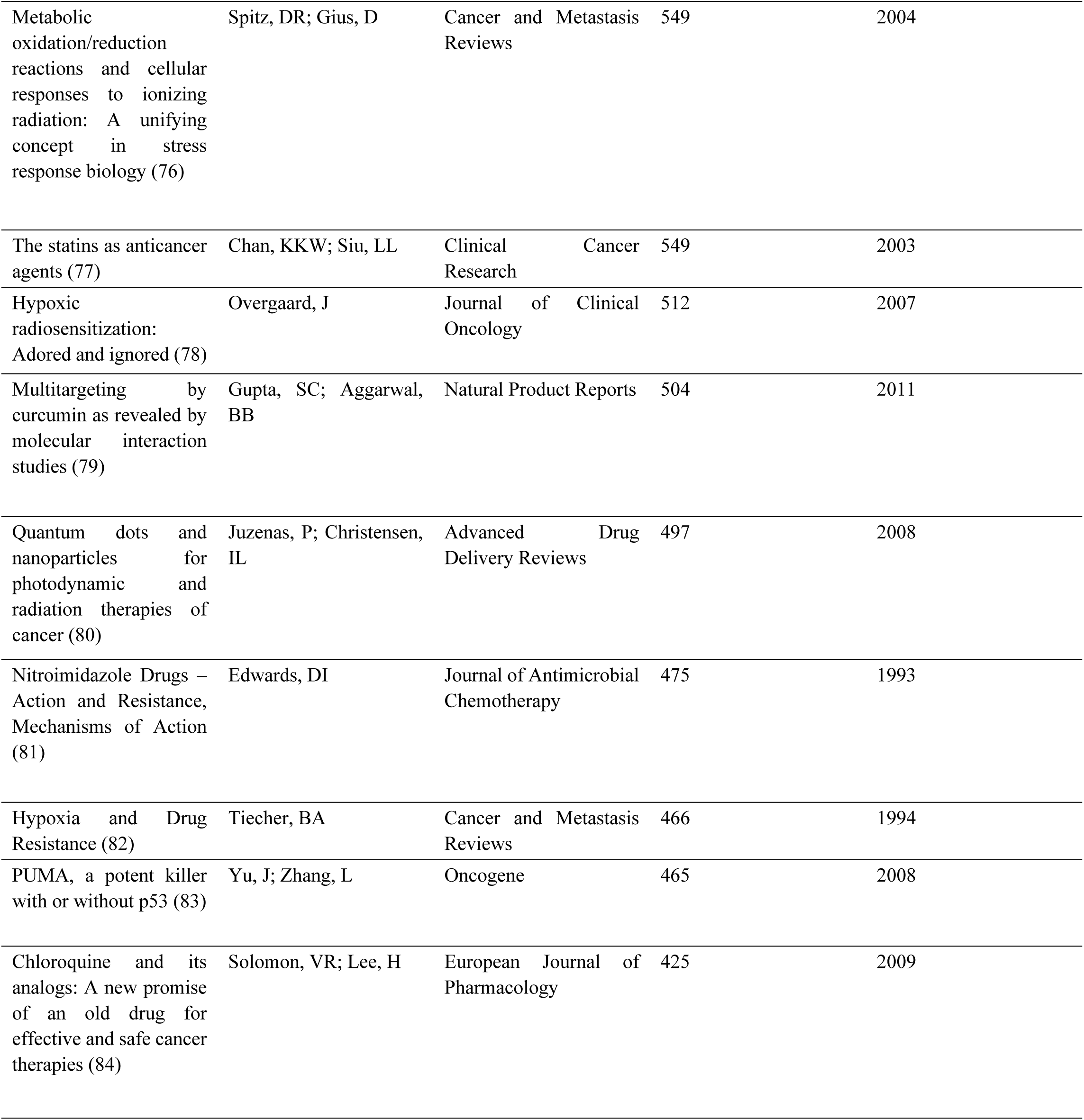
The Top 20 Review Radiosensitizer Publications.

#### 3.4.2 Prominent Authors by Local H-Index and G-Index

The most prominent authors were ranked by local H-index. Lawrence, TS, with a local H-index of 37, was the most prolific and well cited author in this list, particularly within the community of radiosensitizer research (Table 10).

**Table 10:** The Top 10 Authors by Local H-index and G-Index.

| Author | Local H-Index | Local G-Index |
| --- | --- | --- |
| Lawrence, TS | 37 | 61 |
| Brown, JM | 31 | 56 |
| Kinsella, TJ | 27 | 42 |
| Mitchell, JB | 27 | 39 |
| Adams, GE | 26 | 51 |
| Camphausen, K | 25 | 42 |
| Tianfeng, C | 25 | 42 |
| Kim, JH | 23 | 39 |
| Milas, L | 23 | 35 |

Reference year spectroscopy indicates that the years with citation counts that significantly deviated from the 5-year median were 1984, 2000, and 2004. These trends suggest that radiosensitizer research tends to follow a regular pattern of publication.

### 3.5. Keyword and Topic Analysis

Each of the nearly 13,000 articles returned from the radiosensitizer search was categorized based on the co-occurrences of keywords. Initial filtered keyword analysis results from VOS-Viewer yielded a thematic map of concepts with defined clusters of gold nanoparticles, DNA repair, radiosensitivity, and chemotherapy. Thematic maps generated through the Bibliometrix R package were utilized to quantitatively identify trends in key words, as well as the dynamics of key representative topics. The most prominent themes in radiosensitizer research identified through key words were DNA damage, gold nanoparticles, chemotherapy, *in vivo* enhancement, and *in vitro*. The DNA damage topic was supported by the prominent key words of DNA, damage, and strand breaks. The gold nanoparticles cluster contained key words gold nanoparticles, nanoparticles, delivery, oxygen, photodynamic therapy, drug-delivery, dose enhancement, and energy. The chemotherapy cluster contained key words chemotherapy, carcinoma, survival, cisplatin, squamous cell carcinoma, and lung cancer. The *in vivo* cluster contained key words *in vivo*, irradiation, enhancement, mechanism, combination, tumors, and hypoxia. The *in vitro* cluster contained keywords expression, *in vitro*, cell death, activation, inhibition, resistance, and DNA damage. The gold nanoparticles and DNA clusters were the most dynamic themes, while chemotherapy, *in vivo* and *in vitro* clusters were the most developed on the basis of total publication count.

## Discussion

### General Trends

In this work, we analyze both broad and field-specific trends within the 12,701 publications pertaining to radiosensitizers from 1956 to 2024. Overall, the field demonstrated robust growth and diversification, with a large increase in publication count and journal representation with time.

The *International Journal of Radiation Oncology Biology Physics* was the most consistently represented journal in radiosensitizer research. In all subgroup analyses, the most cited studies were published in general radiation and/or oncology journals.

The United States, China and Germany were the most productive countries in radiosensitizer research in terms of total record count. China demonstrated the fastest growth in annual publication count with time, suggesting that it may overtake the United States as the global leader in absolute production of articles in this field. Collaborations in radiosensitization research are the strongest between countries that currently contribute a substantial amount to the total number of publications in the field. Collaboration links are weaker between the top countries that produce radiosensitization research and countries that are emerging in the field. More diverse international collaboration may facilitate the growth of this field.

Global analysis of the most cited primary and review publications suggests that the most universally researched radiosensitizers across all organ subtypes are chemo radiosensitizers, gold nanoparticles, paclitaxel, PARPis, bortezomib, nitroimidazole, chloroquine, and natural compounds such as caffeine and curcumin.

Breast cancer related radiosensitizer research is the fastest growing subfield in radiosensitizer research as characterized by annual record count. Gynecological cancer radiosensitizer research is the slowest growing subfield, potentially due to the historically strong reliance on cisplatin chemo-radiosensitization for cervical cancers, limiting the pursuit of alternative or high efficacy radiosensitizers (85).

### Brain Cancer Radiosensitizers

In the context of brain cancer related radiosensitizer research, chemoradiation investigations were prominent in the 20^th^ century, while articles from the 21^st^ century highlight nanoparticles, ATM inhibitors, mTOR inhibitors and inhibition of PARP. The majority of the most highly cited brain cancer related radiosensitizer publications utilized either syngeneic or xenograft glioma cell lines. The most common syngeneic model utilized was the GL-261 cell line. GL-261 continues to be one of the most widely used syngeneic glioma models in glioma research. However, it has been increasingly identified as less reliable by comparison with newly developed models such as CT-2A or SB-28 (86).

The substantial degree of radioresistance in gliomas and ubiquitous use of adjuvant radiotherapy as a standard of care for high grade gliomas highlights the need for effective radiosensitization to increase the therapeutic receipt and improve patient outcomes(87). As recapitulated in this analysis, the radiosensitizers investigated in pre-clinical studies for gliomas include ATM, mTOR and PARPis. However, radiosensitizers investigated in clinical studies focus primarily on chemoradiotherapies and oxygen mimicking compounds, providing a potential explanation for the numerous pre-clinical studies identified in this analysis as opposed to clinical trials (88). France contributes slightly more to brain – related radiosensitizer research than other subgroups, suggesting either an inherent epidemiological variation, or a coincidental targeted effort.

### Non-Brain Cancer Radiosensitizers

Publications related to lung cancer radiosensitization were predominantly focused on chemoradiation, PARPis, wortmannin, and caffeine. Chemoradiation remains the most utilized radiosensitizer in the treatment of lung cancer (89). Compounds that mimic the electrophilicity of oxygen have the potential to radiosensitize lung cancers due to tendency of these cancers to form hypoxic tumor microenvironments (90). South Korea contributes more to lung cancer radiosensitizer research than radiosensitizer research in general, indicating a specific research focus for the country.

The highly cited study published in *Nature*, “Association of reactive oxygen species and radioresistance in cancer stem cells”, utilized both breast and head and neck cancer cell lines to demonstrate that inhibition of ROS scavengers in cancer stem cells with buthionine sulfoximine may serve as an effective radiosensitizer for a variety of cancers, with direct pre-clinical evidence for breast and head and neck cancer (31). In the field of ENT cancer radiosensitizer research, the most cited publications were chemoradiation focused, except for one study that investigated the role of ß1 integrin signaling in mediating radioresistance (35).

The field of breast cancer radiosensitization research has demonstrated the greatest growth over time, with the most cited publications focused on chemoradiation and gold nanoparticles. Major radiosensitizers relevant for breast cancer also include mTOR inhibitors, ATM inhibitors and PLK-4 inhibitors. However, beyond chemoradiation and precision targeted therapy treatments such as trastuzumab, there is limited substantiation of novel radiosensitizers in clinical trials (91). The most cited studies related to gynecological cancer radiosensitizers focused on gold nanoparticles, suicide gene therapy, anti-tumoral antibiotics and curcumin. However, much of the clinically validated radiosensitizers for gynecological malignancies such as cervical cancers continue to be chemotherapy agents (92).

### Reference Spectroscopy and Keyword Analysis

Reference spectroscopy indicates that in recent years, the field of radiosensitization research has been consistently productive in terms of citations, with minimal deviations in total publication count per year. Keyword and topical analysis identified gold nanoparticles and DNA related mechanistic publications as the most dynamic in relevance over time, while chemoradiation focused studies were more consistently relevant in the field over time. *In vivo* studies tend to be mechanistic in nature and focus substantially on potential combination therapies and hypoxia research. In vitro studies seem to directly study DNA associated mechanisms, cell death, and tumor resistance, and are the oldest most foundational works in the field. Such evidence suggests differential focuses between pre-clinical and clinical studies.

## Conclusion

This bibliometric analysis characterizes the diverse growth and expansion of the field of radiosensitization research – particularly its permeation into cancers of various organ systems, as well as technological and scientific advancements specific to cancers of interest. We present the most highly researched radiosensitizers, as well as the greatest contributing authors, countries, journals and institutions that have defined the field both historically and in the most recent years. Through analysis of the most cited studies in the field, as well as reference spectroscopy and key word analysis, we further identify that the landscape of radiosensitizer research contains a gap in the translation of novel, high efficacy radiosensitizers to clinical practice. For example, although gold nanoparticles have been identified as a high efficacy radiosensitizers for nearly two decades, chemoradiation continues to be the most clinically relevant radiosensitizer for many cancers. The origins and causes of such a gap may be multifaceted but nonetheless demand further evaluation and consideration to ensure improved patient outcomes.

## Declarations

### Ethics Approval and Consent to Participate

Not applicable

### Consent for Publication

All authors consent to the publication of this work.

### Availability of supporting data

Not Applicable

### Author Contributions

Conceptualization, S.A., Y.S.H, and D.J.P.; methodology, S.A. and Y.S.H; software, S.A.; validation, S.A., P.M.H., and Y.S.H..; formal analysis, S.A.; investigation, S.A.; resources, S.D.C. and D.J.P.; data curation, S.A. and P.M.H.; writing—original draft preparation, S.A. and P.M.H; writing—review and editing, S.A, Y.S.H, P.M.H, F.C.L, D.A, A.T, L.U, S.C.E, S.D.C, and D.J.P; visualization, S.A.; supervision, Y.S.H., S.D.C., and D.J.P.; project administration, Y.S.H. and D.J.P. All authors have read and agreed to the published version of the manuscript.

## Appendix A

### Appendix A.1: Additional Search Terms

Keywords for organ specific publications involved a combination of the keywords utilized to gather all radiosensitizers, as well as an ‘AND’ Boolean with another set of keywords describing the organ system of interest. For brain, the additional keyword utilized was ‘brain’. For lung, the additional keyword utilized was ‘lung’. For ENT, the additional keywords utilized were ‘head and neck cancer’, ‘ENT cancer’, ‘oropharyngeal’, ‘nasopharyngeal’, ‘laryngeal’, ‘hypopharyngeal’, ‘oral cancer’, ‘throat cancer’ and ‘sinus cancer’. For GI, the additional keywords utilized were ‘gastrointestinal’, ‘GI’, ‘esophageal’, ‘gastric’, ‘colorectal’, ‘colon’, ‘rectal’, ‘anal’, hepatocellular’, ‘liver’, ‘pancreatic’, ‘pancreas’, ‘bile duct’, ‘gallbladder’, and ‘cholangiocarcinoma’. For gynecology, the additional keywords utilized were ‘gynecologic’, ‘gyn’, ‘cervical’, ‘endometrial’, ‘uterine’, ‘ovarian’, ‘vaginal’ and ‘vulvar’. For Genitourinary, additional keywords utilized were ‘genitourinary’, ‘GU’, ‘prostate’, ‘bladder’, ‘kidney’, ‘renal’, ‘RCC’, ‘urothelial’, ‘testicular’, and ‘penile’. For breast cancer, the additional keywords utilized were ‘breast’, ‘mammary’, ‘ductal’, ‘lobular’, ‘inflammatory’, ‘TNBC’, and ‘triple negative breast cancer’. All keywords within organ subgroups were separated by an ‘OR’ Boolean.

